# Automatic quality control of single-cell and single-nucleus RNA-seq using valiDrops

**DOI:** 10.1101/2023.02.07.526574

**Authors:** Gabija Kavaliauskaite, Jesper Grud Skat Madsen

## Abstract

Single-cell and single-nucleus RNA-sequencing (sxRNA-seq) measures gene expression in individual cells or nuclei, which enables unbiased characterization of cell types and states in tissues. However, the isolation of cells or nuclei for sxRNA-seq can introduce artifacts, such as cell damage and transcript leakage. This can distort biological signals and introduce contamination from debris. Thus, the identification of barcodes containing high-quality cells or nuclei is a critical analytical step in the processing of sxRNA-seq data. Here, we present valiDrops, which is a novel data-adaptive method to identify high-quality barcodes and flag dead cells. In valiDrops, barcodes are initially filtered using data-adaptive thresholding on community-standard quality metrics and subsequently, valiDrops uses a novel clustering-based approach to identify barcodes with biological distinct signals. We benchmark valiDrops and existing methods and find that the biological signals from cell types and states are more distinct, easier to separate and more consistent after filtering by valiDrops. Finally, we show that valiDrops can be used to predict and flag dead cells with high accuracy. This novel classifier can further improve data quality or be used to identify dead cells to interrogate the biology of cell death. Thus, valiDrops is an effective and easy-to-use method to remove barcodes associated with low quality cells or nuclei from sxRNA-seq datasets, thereby improving data quality and biological interpretation. Our method is openly available as an R package at www.github.com/madsen-lab/valiDrops.

## Introduction

The widespread adaptation of single-cell and single-nucleus RNA-sequencing (sxRNA-seq) is producing new and revolutionizing insight into the function of cells and tissues. However, during the isolation of cells or nuclei for sxRNA-seq, they can become apoptotic, stressed, or damaged. The magnitude of these artifacts is protocol-specific and can lead to distortion of the biological signal for example through activation of early-response factors, induction of apoptosis, and transcript leakage (Denisenko et al., 2020; Ilicic et al., 2016; Machado et al., 2021; O’Flanagan et al., 2019). In addition to distorting the biological signal, these processes also lead to the contamination of the solution with debris, such as cell-free ambient RNA (Yang et al., 2020; Young and Behjati, 2020). This problem is exacerbated when processing solid tissues, where harsh methods can be required to release single cells, or when processing nuclei, where cells are lysed, and their contents are released.

Currently, most high-throughput sxRNA-seq methods use either combinatorial indexing (e.g., sci-RNA-seq (Cao et al., 2017), SPLiT-seq (Rosenberg et al., 2018), and scifi (Datlinger et al., 2021)), or are based on droplet emulsions (e.g., Chromium Single Cell Gene Expression (Zheng et al., 2017), Drop-seq (Macosko et al., 2015), inDrop (Klein et al., 2015), or HyDrop (De Rop et al., 2022)). In droplet-based methods, microfluidics is used to process single cells or nuclei in a water-in-oil emulsion, which contains the necessary reagents to synthesize cDNA with droplet-specific barcodes. Debris may be captured together with cells or nuclei adding noise to the biological signal, or debris may be encapsulated into empty droplets creating unwanted signals. Thus, the isolation process can introduce at least three analytical challenges: to separate contaminated empty droplets from cell- or nucleus-containing droplets; and to separate cell- or nucleus-containing droplets with a high signal-to-noise ratio from those with a low signal-to-noise ratio; and to identify droplets containing cells or nuclei, which were not strongly affected by the isolation process.

Several methods have been developed to address the first two challenges; EmptyDrops (Lun et al., 2019) fits a Dirichlet-Multinomial distribution to an estimated ambient RNA and removes droplets, whose expression profile does not significantly differ from the fitted distribution. CB2 (Ni et al., 2020) extends EmptyDrops by introducing a clustering step prior to comparison with the fitted ambient RNA distribution. DIEM (Alvarez et al., 2020) clusters barcodes initially using K-means clustering on principal components and then optimizes the cluster labels using a semi-supervised expectation-maximization algorithm. Finally, DIEM removes contaminated barcodes if they are assigned to debris clusters or have high expression of genes significantly enriched in debris clusters. Both EmptyNN (Yan et al., 2021) and CellBender (Fleming et al., 2019) use deep learning models to identify and remove empty and highly contaminated barcodes. EmptyNN employs positive-unlabeled learning to directly identify which barcodes to remove, while CellBender uses an unsupervised generative model to learn the background RNA profile and recover uncontaminated counts from non-empty droplets. Finally, dropkick (Heiser et al., 2021) automatically labels barcodes as informative or non-informative based on the number of detected genes and then refines labels using a logistic regression model with elastic net regularization.

Except for dropkick, all the above methods are dependent on a threshold that separates cell- or nucleus-free barcodes from putative cell- or nucleus-containing barcodes. Furthermore, except for CellBender, DIEM, and dropkick, the above method requires additional filtering to remove barcodes, which contain a low-quality cell or nucleus. Commonly, this is done by thresholding on the fraction of unique molecular identifiers (UMIs) derived from mitochondrial genes, the total number of detected genes, and the total number of UMIs (Luecken and Theis, 2019).

Here, we present valiDrops; an automated method for identifying cell- or nucleus-containing barcodes with a high signal-to-noise ratio and low signal distortion. Our method is open source and available as an R package at www.github.com/madsen-lab/valiDrops. We showcase how each step in valiDrops improves the data quality and we extensively benchmark valiDrops and existing methods using 47 real samples from five different studies to show that valiDrops has the best performance for both single-cell and single-nucleus RNA-seq data. Finally, we demonstrate how valiDrops, unlike existing methods, is also able to detect and flag dead cells to further increase data quality or to enable interrogating of the biological processes leading to cell death.

## Results

### Overview of methods and the benchmarking strategy

valiDrops takes as input an unfiltered feature-by-barcode matrix, which can be produced by all common alignment and count methods, for example CellRanger (Zheng et al., 2017), STARsolo (Kaminow et al., 2021), kallisto | bustools (Melsted et al., 2021), and alevin (Srivastava et al., 2019), and sequentially removes barcodes of low quality (Figure 1A). Some of the quality metrics used in valiDrops for barcode filtering, such as the total number of unique molecular identifiers (UMIs), are already extensively used in the field, but often require user input to set thresholds. In contrast, valiDrops automatically detects thresholds, such as the threshold defining the ‘ambient plateau’ and remove barcodes with a low number of UMIs, which are likely to contain only ambient signals. Next, valiDrops uses stepwise linear regression to infer the relationship between the number of detected features and the number of UMIs and automatically removes outliers. It uses knee-point detection to automatically detect an appropriate threshold for filtering based on the fraction of UMIs derived from mitochondrial genes, and it fits a normal distribution to the fraction of UMIs derived from protein-coding genes and sets a threshold using the fitted mean and standard deviation. In addition to these commonly used quality metrics, valiDrops uses a novel filtering approach based on differentially expressed genes. Barcodes are grouped into both high-resolution and low-resolution clusters using a combination of graph clustering and a data-adaptive method for selecting clustering resolutions. The barcodes in each high-resolution cluster are then compared to all other barcodes, which do not belong to the same low-resolution cluster, and any high-resolution clusters that do not show significant differences in gene expression compared to the other barcodes are removed. This process helps to remove to remove droplets containing a cell or nucleus with a low signal-to-noise ratio. Finally, valiDrops has the option to predict and flag putative dead cells. To detect dead cells, valiDrops initially labels dead cells using a simple heuristic based on the fraction of UMIs assigned to mitochondrial, ribosomal, and protein-coding genes. Next, valiDrops refines these noisy labels using ridge regression and an adaptive resampling strategy.

**Figure 1:**
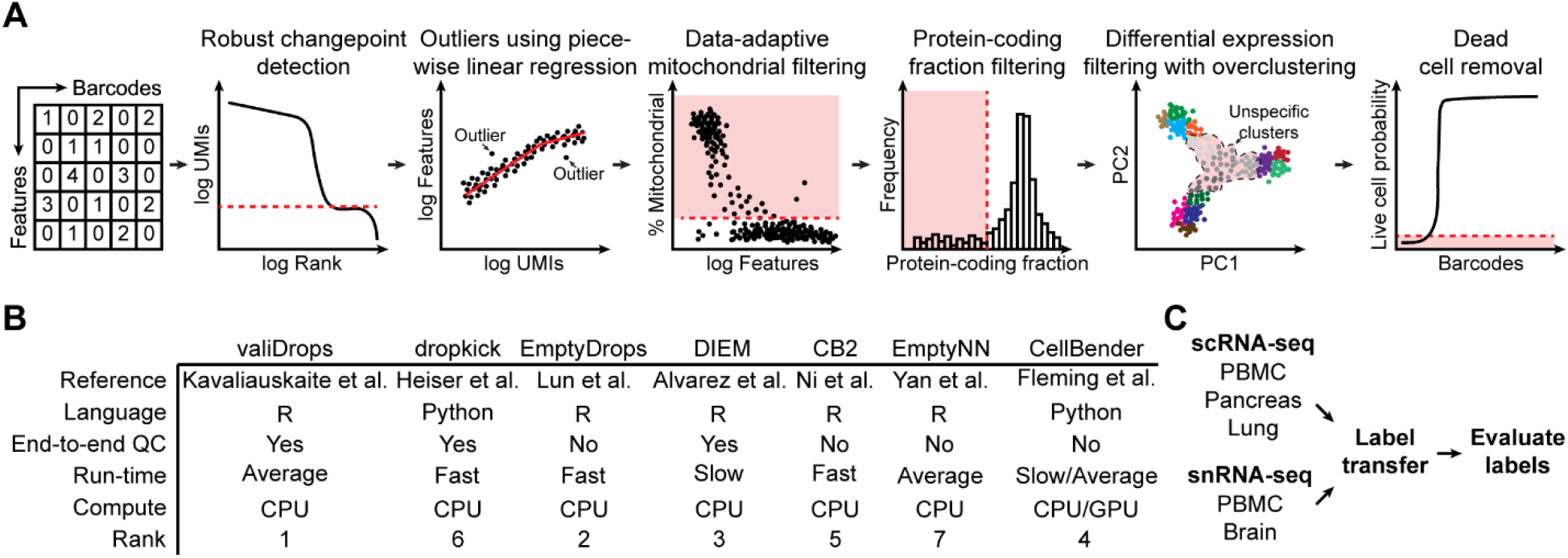
Overview of valiDrops and benchmarking methods and results. **A)** Illustration of the major steps in valiDrops. **B)** Overview of existing methods in the field, specifying the programming language, whether they require post-filtering, an estimate of their run time, the compute backend they use and the rank they achieved in our benchmarking. * CellBender (Fleming et al., 2019) is significantly faster if run using a GPU. **C)** Overview of the benchmarking approach and datasets used to valiDrops and benchmark to existing tools. The indicated datasets were quality filtered using valiDrops or alternative methods. Azimuth (Hao et al., 2021) was used to transfer labels from reference atlases and the resulting transferred labels were evaluated in terms of homogeneity and separability.

Here, we benchmarked valiDrops and six other existing tools and find that valiDrops has the best performance and an average run-time (Figure 1B). It is an open problem how to benchmark barcode filtering methods since there are no ground truth datasets available. Here, we devised a strategy based on assessing the quality of transferred labels after barcode filtering (Figure 1C). For each tool, we filtered barcodes across 47 single-cell or single-nucleus RNA-seq samples from five different datasets and subsequently used Azimuth (Hao et al., 2021) to transfer cell type labels. Then, we assessed the quality of the identified barcodes by evaluating labeling metrics, conservation of label hierarchies, clustering of labels, and intra-label transcriptional entropy. Additionally, we integrated samples from the same dataset using Seurat (Hao et al., 2021), and assessed the extent of batch effects by evaluating the conversation of biological signals, separation of batch labels, and similarity of cell type labels using both cluster-dependent and cluster-independent metrics.

### valiDrops sequentially removes low-quality barcodes

To evaluate how each stage in valiDrops improves data quality, we analyzed a single PBMC dataset from 10X Genomics containing 5,000 PBMCs (10X, 2019) and transferred labels using Azimuth (Hao et al., 2021) to barcode passing each of three stages in valiDrops. In stage 1, valiDrops removed barcodes with a low number of UMIs. In stage 2, valiDrops removed barcodes using common quality metrics, such as the number of UMIs derived from mitochondrial genes and the relationship between the number of UMIs and number of detected genes. In stage 3, valiDrops removed barcodes without distinct biological signals using a data-adaptive clustering approach and differential expression testing. After stage 1, most transferred cell type labels are well separated in the UMAP space (Figure 2A, left panel). However, major clusters of monocytes and T-cells are connected by putative erythrocytes, which are also found in several places across the embedded space (black arrows). After stage 2, the cell types are clustered more tightly. Most erythrocytes connecting major clusters of monocytes and T-cells have been removed, although a tail in both clusters is retained (Figure 2A, middle panel). Finally, after stage 3, the cell types are more well-separated in the UMAP space, and the tails of erythrocytes are removed leaving one distinct cluster of putative erythrocytes (Figure 2A, right panel).

**Figure 2:**
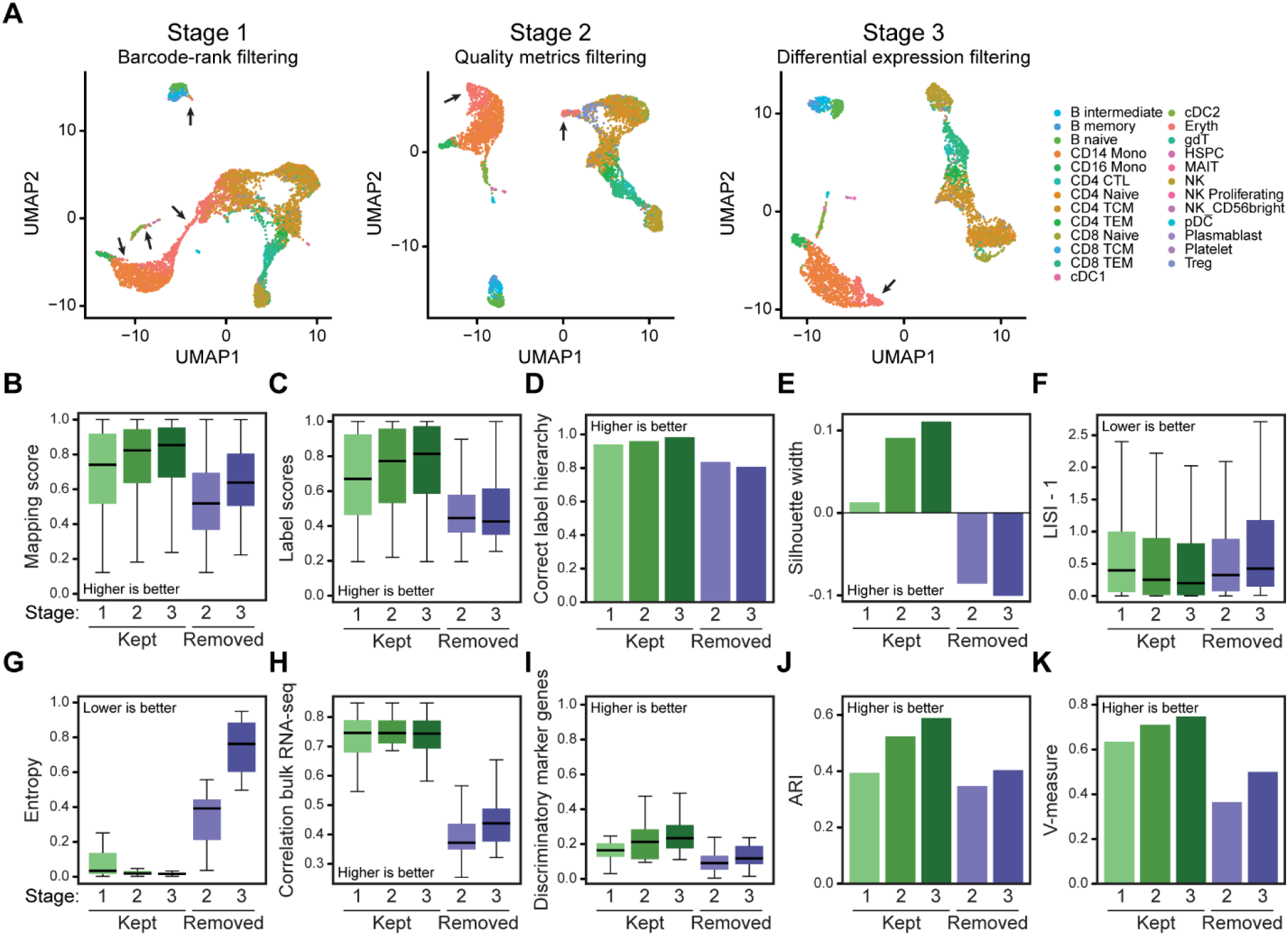
Quality filtering using valiDrops improves data quality. **A)** Barcodes that pass filtering on total UMI counts (stage 1), filtering on quality metrics (stage 2) and on expression-based metrics (stage 3) from a reference PBMC sample from 10X Genomics (10X, 2019) were embedded in UMAP space using Seurat (Hao et al., 2021) based on 10 PCs calculated from 2000 highly variable genes. Labels were transferred using Azimuth (Hao et al., 2021) with its associated PBMC reference atlas. The arrows point to erythrocytes. **B-K)** Metrics for barcodes that pass filtering in stage 1, 2 or 3, as well as for barcodes which are removed between either stages 1 and 2 or between stages 2 and 3: **B)** Boxplot of mapping scores calculated by Azimuth during label transfer. **C)** Boxplot of label scores calculated by Azimuth during label transfer. **D)** The fraction of labels which have the correct label hierarchy **E)** The average silhouette width across transferred labels in the first 10 PCs. **F)** Median local inverse Simpson’s index (LISI) across transferred labels in the first 10 PCs. **G)** Entropy of the top 200 mostly highly expressed genes in each label class. **H)** Boxplot of median Spearman’s correlation coefficient between pseudo-bulk expression levels and the expression levels in RNA-seq from sorted cells of the same cell type (Monaco et al., 2019). **I)** The fraction of marker genes in a label class, defined as differentially expressed genes (FDR ≤ 0.05, log fold change > 0) compared to all other barcodes, with an AUC higher than 0.9. **J)** Barplot showing the highest ARI across multiple resolutions of Louvain-based clusters and transferred labels (see Methods). **K)** Barplot showing the highest V-measure across multiple resolutions of Louvain-based clusters and transferred labels (see Methods).

To quantify the extent to which each of the stages improved the quality of the dataset, we used a large and diverse panel of measures related to labeling, cell type separability, transcriptome similarity, and cell type clusterability (i.e., the precision at which clustering can separate cell type labels) (Figure 2B-K) (see Methods). We find that labeling scores and the accuracy of the label hierarchy improved for each stage and that the removed barcodes have much lower scores than the retained barcodes (Figure 2B-D). To determine the accuracy of the label hierarchy, we labeled the barcodes with both coarse-grained and fine-grained labels and asked how large a fraction of barcodes was assigned to a fine-grained label, which was a child of the assigned coarse-grained label. We found a similar trend across all metrics; cluster separability increased across filtering stages and removed barcodes have low separability (Figure 2E-F). Transcriptomic similarity, correlation to bulk RNA-seq expression, and the fraction of markers that were discriminatory also increased across filtering stages (Figure 2G-I), and so did the ability to cluster barcodes into cell types (Figure 2J-K). Collectively, this strongly indicates that valiDrops filtering improves the quality of the dataset.

### valiDrops compares favorably to existing tools

To compare valiDrops to existing methods, we ran all methods, except CellBender, using default parameters. CellBender does not have a default parameter set, as it requires users to specific the number of expected cells, as well as the number of total cells. In this benchmark, we derived these numbers from the automatically determined thresholds set by valiDrops. Finally, one of the key distinguishing features of valiDrops is that it is automated, does not require post-filtering, and does not require user input. In contrast, EmptyDrops, scCB2, and EmptyNN all require post-filtering to remove barcodes containing low quality cells or nuclei. To ensure that these tools did not artificially underperform due to a lack of post-filtering, we applied miQC (Hippen et al., 2021) to automatically filter barcodes based on mitochondrial content for the single-cell RNA-seq samples and set a data-driven mitochondrial threshold at three times the median absolute deviation above the median for single-nucleus RNA-seq samples.

Initially, we evaluated a single PBMC dataset from 10X Genomics containing 8,000 PBMCs (10X, 2017) (Figure 3A). All methods detected approximately 8,000 high-quality barcodes, except dropkick and EmptyNN, both of which detected more than 9,000 barcodes (Figure 3B). Across the labeling metrics, valiDrops achieved the highest mapping score, but scores were overall similar (Figures 3C-E). However, barcodes identified by valiDrops had the highest transcriptome similarity within cell types, as measured by entropy (Figure 3F). For cell type cohesion and clusterability (see Methods), valiDrops achieved the best scores for the silhouette width, for the local inverse Simpson’s index (LISI) (Figures 3G-H) and for all clustering metrics (Figures 3I-K). Thus, in this dataset, valiDrops exhibited the best performance across all metrics.

**Figure 3:**
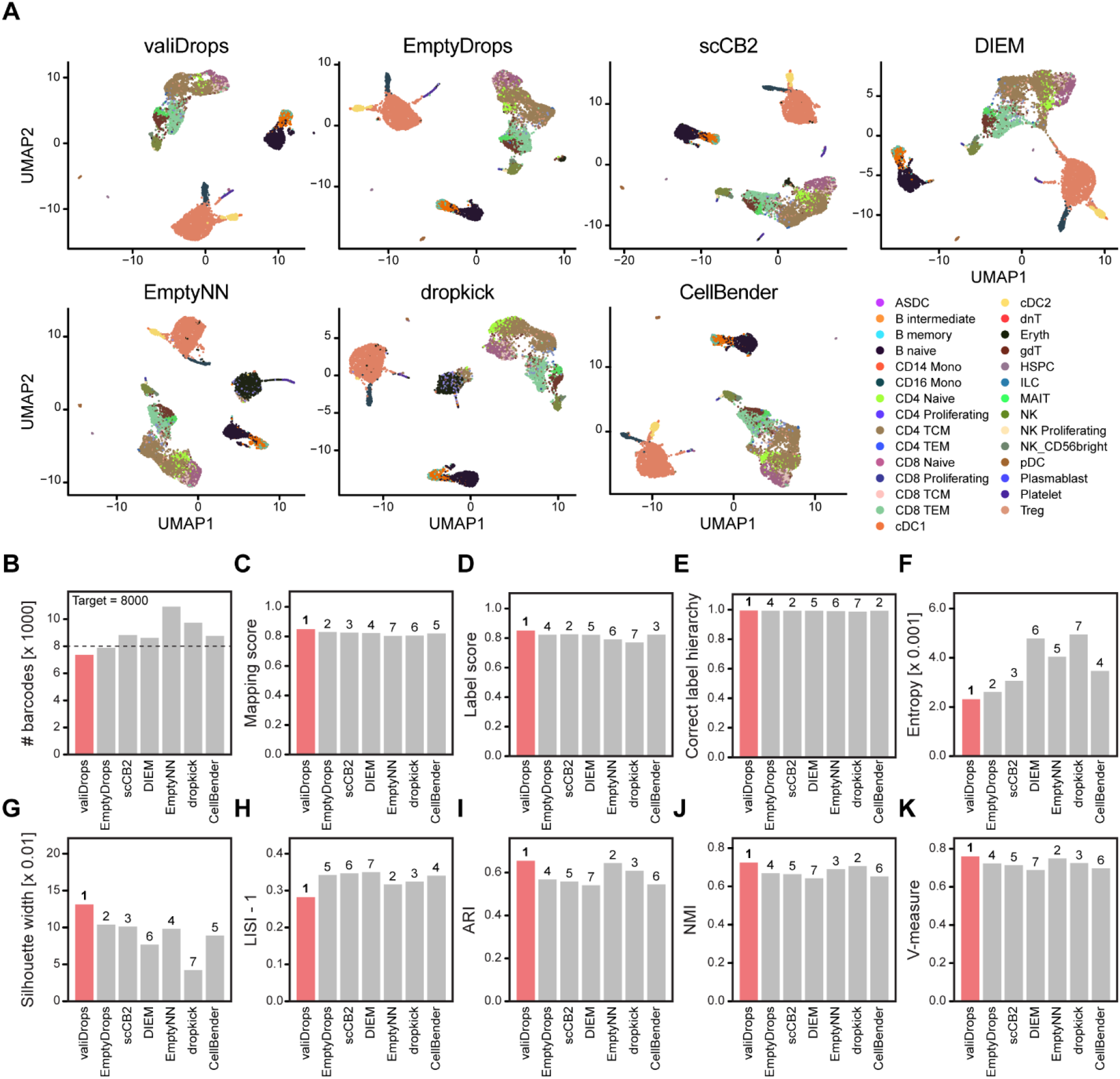
Comparison of valiDrops and alternative quality control methods on PBMC data. **A)** Barcodes that pass filtering by valiDrops, EmptyDrops (Lun et al., 2019), DIEM (Alvarez et al., 2020), scCB2 (Ni et al., 2020), dropkick (Heiser et al., 2021), CellBender (Fleming et al., 2019), or EmptyNN (Yan et al., 2021) from a reference PBMC samples from 10X Genomics (10X, 2017) were embedded in UMAP space using Seurat (Hao et al., 2021) based on 10 PCs calculated from 2,000 HVGs. Labels were transferred using Azimuth (Hao et al., 2021). **B)** The number of barcodes that pass filtering in each method. Vertical line; the experimentally targeted number of cells. **C-K)** Metrics for barcodes filtered as in A. The numbers indicate the rank of the method, where 1 is the best and 7 is the worst and valiDrops is highlighted in red. **C)** Average label transfer mapping score calculated by Azimuth. **D)** Average label transfer score calculated by Azimuth. **E)** The fraction of labels that have the correct label hierarchy (see Methods). **F)** Entropy of top 200 mostly highly expressed genes in each label class. **G)** Average silhouette width across transferred labels in the first 10 PCs. **H)** Median local inverse Simpson’s index (LISI) across transferred labels in the first 10 PCs. **I-K)** Max ARI (I), NMI (J), or V-measure (K) across multiple resolutions of Louvain-based clusters and transferred labels (see Methods).

We expanded this benchmark by analyzing a total of 47 sxRNA-seq samples from five datasets. Each method was ranked based on the set of nine metrics (the overall rank), as well as on subsets of metrics that were used to evaluate label mapping, expression similarity, label cohesion, and label clusterability (Supplemental Table S1). In addition, we integrated the samples from each dataset using Seurat and evaluated the integration in terms of marker gene conservation, cross-sample gene-rank correlation, integration metrics, and biological conservation metrics (Supplemental Table S2). Each method was ranked based on the full set of 12 metrics (the overall rank) and on subsets of metrics, which were used to evaluate expression similarity, label cohesion, label clusterability, batch cohesion, and batch clusterability (see Methods). For all five datasets and integration tasks, valiDrops achieved the best overall rank (Figure 4A). For filtering individual samples, valiDrops achieved the best overall rank for 35 of the 47 samples, rank in the top 2 for 44 of the 47 samples and never ranks worse than 4 and this high performance was consistent across all metrics (Figure 4B). For batch integration after filtering, valiDrops achieved the best overall rank for all tasks, but notably, did not consistently achieve a high rank for expression similarity. However, looking at the individual metrics that compose this rank, we found that all methods perform approximately similarly (Figures 4C-D). Thus, valiDrops provides better barcode filtering compared to existing methods.

**Figure 4:**
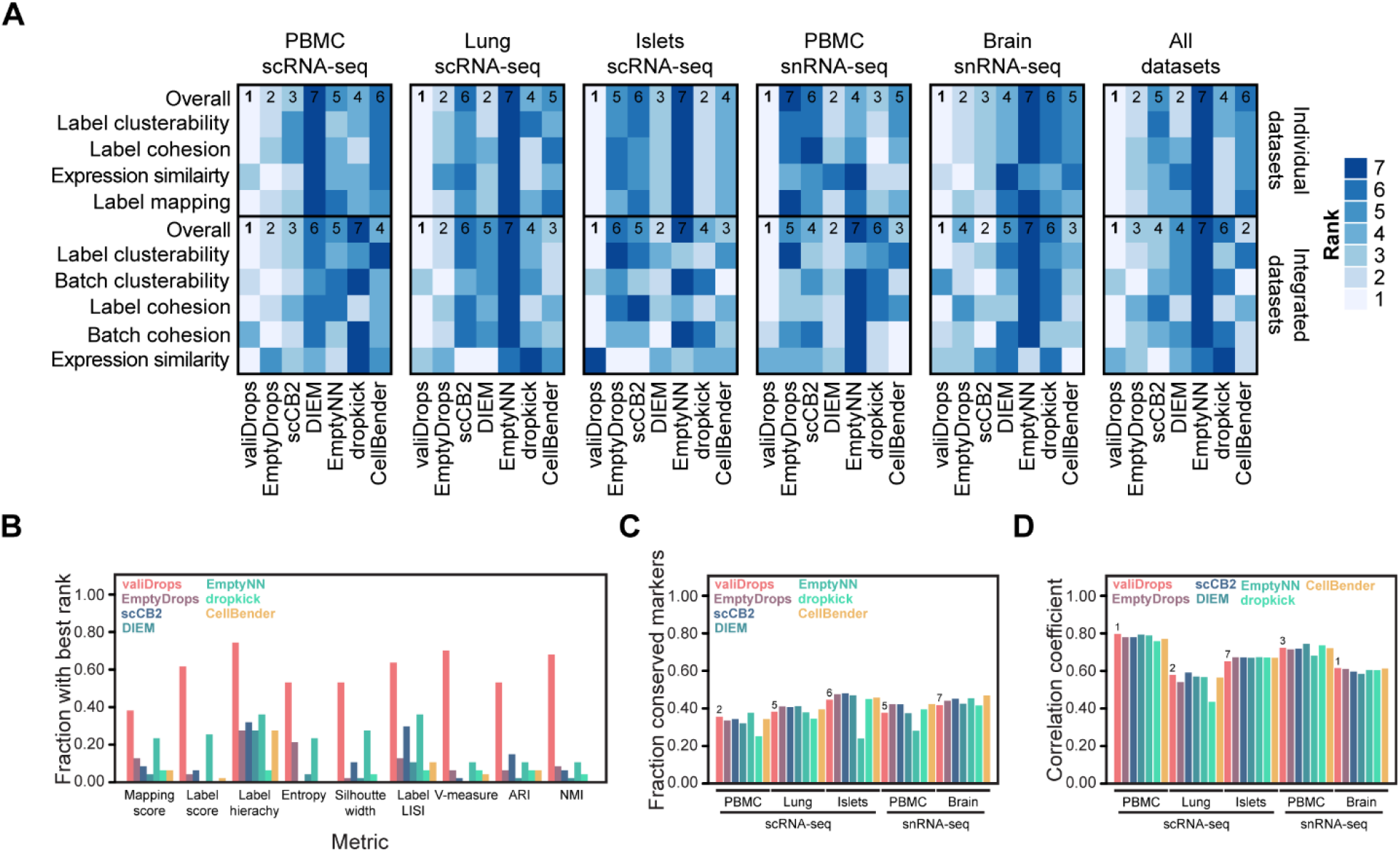
Benchmarking valiDrops and alternative quality control methods. Barcodes were quality filtered using valiDrops, EmptyDrops (Lun et al., 2019), DIEM (Alvarez et al., 2020), scCB2 (Ni et al., 2020), dropkick (Heiser et al., 2021), CellBender (Fleming et al., 2019), or EmptyNN (Yan et al., 2021) and labels were transferred with Azimuth (Hao et al., 2021) with associated reference atlases. The value in each table is the overall rank of the method based on individual dataset metrics, where lower is better (see Methods of a detailed description of the metrics). For the integration rank, each sample was quality controlled individually and integrated using the RPCA method in Seurat (Hao et al., 2021) with default parameters. The value in each table is the overall rank of the method based on integration metrics, where lower is better (see Methods of a detailed description of the metrics). **A)** Heatmap for ranks for all datasets, as well as an overall rank across all datasets. **B)** Barplot showing the fraction of datasets that achieved the best rank for each metric and for each method. **C)** Barplot showing the fraction of conserved marker genes across the five integration datasets for each method. **D)** Barplot showing the median Spearman’s rank coefficient of correlation between pseudo-bulk expression levels per class label per dataset across the five integration datasets for each method.

### valiDrops can detect dead cells with high accuracy

Recent literature suggests that a subset of dead cells can pass regular quality control and confound the transcriptomic signatures of cell types or states in sxRNA-seq dataset (O’Flanagan et al., 2019; Ordonez-Rueda et al., 2020). This issue may be especially prevalent in certain diseases associated with the induction of cell death, or when storing tissues for later processing, where cell types within the tissue may have a differential sensitivity to the storage conditions.

To overcome such potential confounding, we designed an optional module in valiDrops to predict dead and live cells. To create this module, we leveraged two datasets with ground truth, where the authors had induced cell death, sorted cells into live and dead populations, and performed scRNA-seq on both groups (O’Flanagan et al., 2019; Ordonez-Rueda et al., 2020). Based on these labels, we created a module with three steps. Initially, every cell is assigned a score based on arcsine-transformed fractions of UMIs derived from mitochondrial genes, ribosomal genes, and coding genes, as well as the standardized number of total UMIs and detected genes. Second, the cells are labeled using a data-adaptive thresholding approach to separate the cells into a group enriched for dead cells, and a mixed group containing both live and dead cells. Finally, the labels are optimized using ridge regression and a modified version of adaptive resampling (Yang et al., 2019). In the two datasets, initial labels made by valiDrops were weak predictors of the true class, whereas the labels optimized using adaptive resampling and ridge regression were highly accurate (Matthews Correlation Coefficient (MCC) ≥ 0.86), and significantly better than baseline models using logistic regression on the initial labels (Figures 5A-B). To evaluate the sensitivity of valiDrops, we used a stratified subsampling approach to approximate the distribution of initial scores but reduce the number of true dead cells by between 10 – 50%. In one dataset, valiDrops maintained a median MCC above 0.9 across the subsampled datasets, while in the other, the MCC decreased to a median of approximately 0.66 after the removal of 50% of the truly dead cells. Although, the MCC decreases, this still represented a strong predictive performance (Figure 5C). The difference in MCC between the datasets was likely explained by the absolute numbers of truly dead cells present after subsampling, as the dataset with the lowest MCC only contained a median of 4 truly dead cells, whereas the dataset with the highest MCC contained a median of 50 truly dead cells. The decrease in MCC observed at small numbers of truly dead cells was driven by a decrease in the ability to correctly label dead cells (Figure 5D), not in the ability to correctly label live cells (Figure 5E). Reassuringly, this highlight that valiDrops does not spuriously flag live cells for removal.

**Figure 5:**
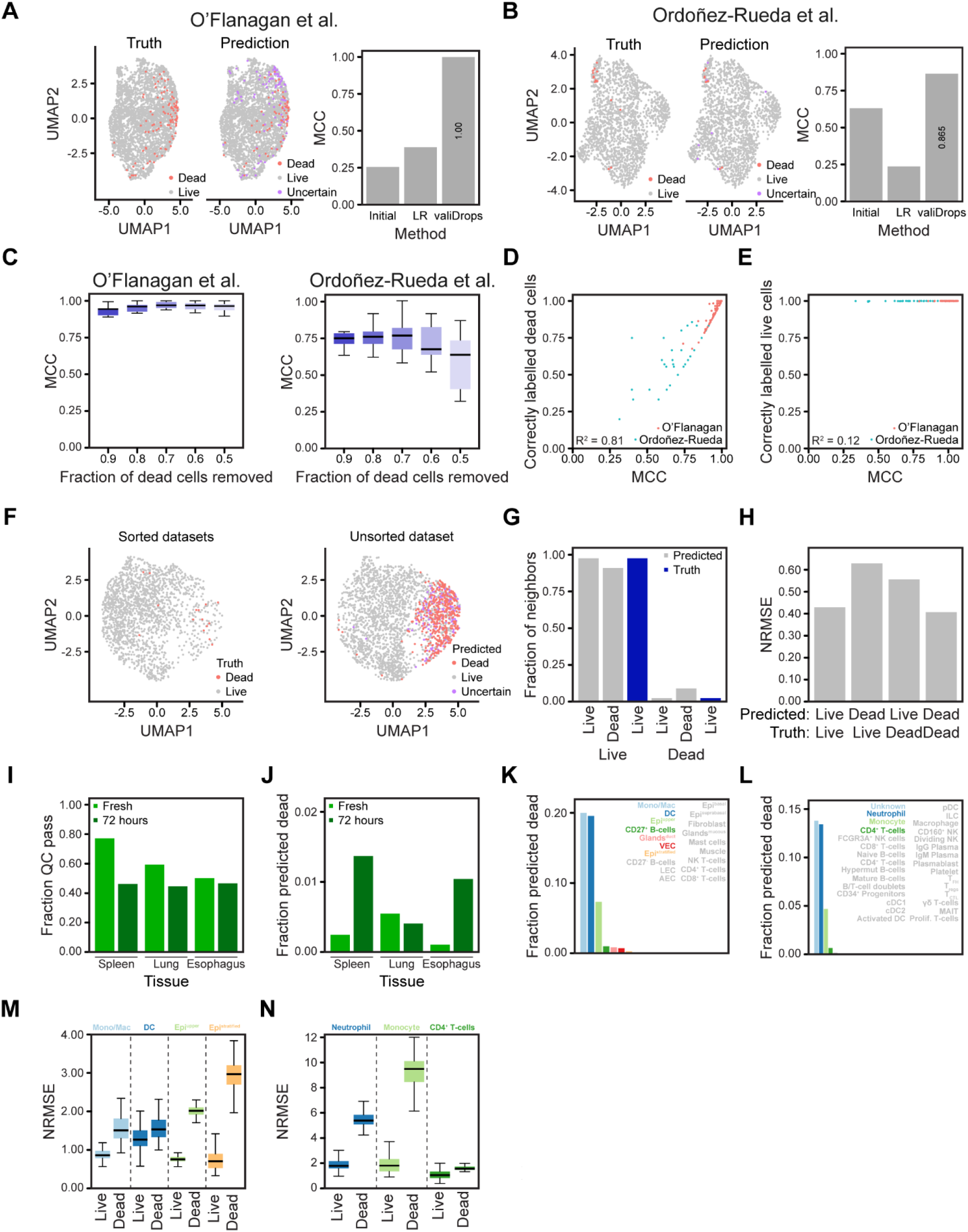
Identification of dead cells using valiDrops. **A-B)** Sorted dead and sorted live cells from the O’Flanagan datasets (O’Flanagan et al., 2019) and the Ordoñez-Rueda datasets (Ordonez-Rueda et al., 2020) were quality filtered using valiDrops and combined before dead cells were predicted. Barcodes that pass quality control were embedded in UMAP space (left; true labels, right; predicted labels) using Seurat (Hao et al., 2021) based on 20 PCs calculated from 500 HVGs. The barplots show the Matthew’s correlation coefficient (MCC) for the initial labels, for a baseline model using logistic regression on the initial labels (LR) and for valiDrops. **C)** Boxplots showing MCC for dead cell prediction in the indicated dataset after removal of the indicated fraction of truly dead cells. **D)** Scatterplot of the MCC and the fraction of truly dead cells that were correctly labelled. **E)** Scatterplot of the MCC and the fraction of truly live cells that were correctly labelled. **F)** UMAP embedding (left; sorted datasets with true labels, right; unsorted datasets with predicted labels) using Seurat (Hao et al., 2021) based on 10 PCs calculated from 100 HVGs of an unsorted sample from the Ordoñez-Rueda dataset (Ordonez-Rueda et al., 2020) after quality filtered and dead cell prediction. **G)** The fraction of ten nearest neighbors derived from truly live (first three bars) or truly dead (last three bars) computed on the 20 first PCs calculated from 100 HVGs for predicted live, predicted dead or truly live cells. **H)** The normalized root-mean-square-error (NRMSE) between pseudo-bulk expression levels for top 100 most highly expressed genes in truly live cells and either predicted live or predicted dead cells, and between the pseudo-bulk expression levels for top 100 most highly expressed genes in in truly dead cells and either predicted live or predicted dead cells. **I)** Barplot of the fraction of barcodes that pass filtering for low total UMI counts (stage 1) that pass final quality filtering for fresh samples and samples stored for 72 hours from the spleen, the lungs, and the esophagus (Madissoon et al., 2019). **J)** The fraction of barcodes that pass final quality control that are predicted to be dead. **K-L)** The fraction of predicted dead cells by cell type labels (K; esophagus, L; spleen). Cell types with more than 1% dead cells were colored. **M-N)** Boxplot across subsamples showing NRMSE between pseudo-bulk expression levels for top 100 most highly expressed genes in fresh cells and either predicted live or predicted dead cells stored for 72 hours for the indicated cell types (M; esophagus, N; spleen). Cell types with at least 1% of the cells, and at least a total of five predicted dead cells after storage for 72 hours, were included in the analysis.

To test the module on unseen data, we processed and predicted dead cells in an additional sample included in one of the ground truth datasets (Ordonez-Rueda et al., 2020), where cell death had been induced, but the cells had not been sorted. To evaluate the predicted labels, we integrated the unsorted and sorted datasets. The predicted dead cells in the unsorted dataset are more often neighbors to truly dead cells than to truly live cells in the integrated reduced dimensional space. Similarly, the predicted live cells are more often neighbors to truly live cells than to truly dead cells (Figures 5F-G). Comparison of the pseudo-bulk transcriptomic profile of predicted live and predicted dead cells to that of truly live and truly dead cells revealed that the predicted live cells are transcriptionally more similar to truly live cells, while predicted dead cells are more similar to truly dead cells (Figure 5H). This suggests that *in silico* dead cell labeling can rescue samples that are contaminated with dead cells or serve as a basis for studying mechanisms of cell death.

To test the module in more biologically relevant systems, we re-analyzed data from a study that assessed the effects of cold storage of up to 72 hours on healthy human spleen, esophagus, and lungs (Madissoon et al., 2019). The original authors showed using TUNEL staining that on the tissue level especially the spleen and the esophagus had increased numbers of dead cells after 72 hours of cold storage. The spleen and lung samples were processed with a dead cell removal kit, and all three tissues were analyzed by single-cell RNA-seq. In the initial quality control using valiDrops, we found that a decreased fraction of barcodes passed quality control after 72 hours of storage, and that this was especially pronounced in the spleen (Figure 5I). This observation was also reported in the original paper based on manual quality control. Across the datasets, cell labeling by valiDrops predicts low numbers of dead cells (Figure 5J), consistent with the esophagus samples having high viability and the lung and spleen samples being filtered using a dead cell removal kit prior to sample preparation. However, valiDrops did predict an increased number of dead cells in the spleen and the esophagus samples after 72 hours consistent with the TUNEL staining. The fraction and the total number of predicted dead cells vary across cell types (Figures 5K-L) suggesting that different cell types have different susceptibilities to dying during cold storage. In cell types marked by high rates of predicted dead cells, the cells predicted to be live are transcriptionally more similar to cells from the same cell type in the fresh samples compared to cells predicted to be dead (Figures 5M-N). Taken together, these results show how valiDrops can accurately identify dead cells from complex tissues and that removing these cells restores the accuracy of aggregated transcriptomic profiles.

## Discussion

Quality control and filtering are the most important preprocessing steps in analyzing sxRNA-seq datasets. The most widely used workflows rely on tools to identify barcodes having transcriptomic profiles that do not resemble the aggregated signal from barcodes with low coverage. These barcodes are then filtered to define high-quality barcodes based on global and user-defined thresholds for the number of detected genes, the UMI count, and/or the fraction of UMIs derived from mitochondrial genes. The individual researcher can bias this process, it can be very time-consuming, and there can be identification biases against the most prevalent cell types whose transcriptome has the strongest resemblance to the ambient profile. To overcome these issues, we have developed valiDrops, which is as an automated software that identifies high-quality barcodes from raw sxRNA-seq count matrices.

In valiDrops, barcodes are automatically filtered by using data-adaptive thresholds on the number of detected genes, the number of UMIs, their association to each other, the fraction of UMIs derived from mitochondrial genes, and the fraction of UMIs derived from coding genes. Next, valiDrops used an overclustering-based approach to identify small groups of barcodes that have a distinct signal compared to the other barcodes passing initial filtering. Thus, unlike existing method, valiDrops does not rely on comparison between high-quality barcodes and low-coverage barcodes, thereby avoiding biases from differences in how cell types contribute to the profile of low-coverage barcodes. The overclustering-based approach uses a combination of clustering resolutions to ensure that barcodes containing cells or nuclei from the same or closely related cell types and states are not compared, thereby avoiding potential biases against for example differentiating cells. However, since valiDrops looks for distinct signals, valiDrops can only accurately identify high-quality droplets in datasets with biological heterogeneity. Therefore, we do not advise users to apply the quality control and filtering module of valiDrops to datasets derived from for example a single, pure cell line. In complex samples, such as tissue or whole organisms, valiDrops is a valuable method for the automatic and unbiased identification of high-quality barcodes.

Prediction of dead cells by valiDrops has the potential to improve data quality by removing technical biases and to unlock the study of cell death-inducing mechanisms, which is relevant in multiple diseases ranging from cancers to metabolic diseases. In valiDrops, dead cells are initially labelled based on the fraction of UMIs derived from mitochondrially-encoded genes. Due to this strategy, valiDrops is not capable of predicting dead cells from single-nucleus RNA-seq data. To assess the ability valiDrops to improve data quality when faced with dead cells, valiDrops was used to predict dead cells samples with low or high rates of dead cells prior to dead cell removal and single-cell RNA-seq. Consistently, valiDrops detected more dead cells in samples with a priori high rates of cell death, and computational removal of the predicted dead cells improved the transcriptomic similarity between the sample in question and control samples with low rates of dead cells. The sensitivity of dead cell prediction in valiDrops is high, as valiDrops achieves a median MCC ≥ 0.75 in samples with as little as 0.1% dead cells. Below this threshold, valiDrops increasingly misclassifies dead cells as live, but not live cells as dead. Thus, valiDrops does not spuriously remove truly live cells even in the absence of truly dead cells.

valiDrops is available as an R package that is available from GitHub (www.github.com/madsen-lab/valiDrops)and requires a single line of code to automatically identify barcodes containing high-quality nuclei or (live) cells.

## Methods

### Quality filtering and dead cell prediction with valiDrops

The first stage in valiDrops is to remove lowly sequenced barcodes that likely represent barcodes primarily containing ambient RNA. The log-transformed total number of UMIs as a function of the log-transformed rank in order of decreasing total UMIs (which is the basis of the so-called Barcode Rank plot) is smoothened using the rolling mean with a bin size is defined using Rice rule 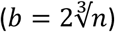. Piece-wise linear regression is fitted to the smoothened curve using segmented. In the default settings, models with 2 – 5 breakpoints are fitted. For each model, the root-mean-square-error (RMSE) of the fit is calculated and the smallest number of breakpoints that have an error with 1.5 times the smallest RMSE is selected. The angle between all piecewise linear curves is calculated and the breakpoint with the smallest angle is selected as the threshold for filtering lowly sequenced barcodes.

The second stage in valiDrops is to collect and filter on common quality metrics. To collect quality metrics, valiDrops initially automatically finds and calculates the fraction of UMIs associated with mitochondrially-encoded genes (mitochondrial fraction), protein-coding genes (coding fraction), and genes encoding ribosomal genes (ribosomal fraction), as well as the log-transformed total number of UMIs and the log-transformed total number of detected genes. This automatic detection works for most common model organisms (human, mouse, rat, fly, worm, and zebrafish) and common gene annotations (Ensembl, Entrez, HGNC, MGI, and gene symbols), but can be overwritten by users studying other organisms or using other gene annotations. Optionally, valiDrops can calculate the fraction of UMIs associated with exons (or introns) that has been shown to be a valuable additional metric for quality filtering (Muskovic and Powell, 2021). To filter on the mitochondrial fraction, valiDrops fits a mixture of two normal distributions to the log-transformed total feature count using mixtools (Benaglia et al., 2009) to identify a group of high-coverage barcodes that are probably high-quality barcodes. For each putative threshold between the median mitochondrial fraction in high-coverage barcodes and one in increments of 0.001, the number of high-coverage barcodes passing the filter is calculated and the final threshold is selected by finding the knee point of the curve using inflection. In rare edge cases with generally high mitochondrial fractions, this approach identifies thresholds above 0.3, which is not biologically plausible (Osorio and Cai, 2021). In these cases, valiDrops uses piece-wise linear regression between the mitochondrial fraction and the log-transformed number of detected genes to set a stricter threshold. To filter on the coding fraction (and optionally the exon or intron fraction), valiDrops calculates the median of the fraction as well as S_n_, which is a more efficient alternative robust scale estimator than the median absolute deviation (Rousseeuw and Croux, 1993). Barcodes more than three times S_n_ above or below the median are removed. Finally, to filter the relationship between the log-transformed total number of UMIs and the number of detected genes, valiDrops fits a piece-wise linear regression model with three break-points, and for each barcode calculates the residuals. Barcodes more than five times S_n_ above or below the median residual are removed.

The third stage in valiDrops is to collect and filter on expression-based metrics. First, 5,000 highly variable genes (HVGs) are selected using scry (Street, 2020), which are then decomposed using singular value decomposition with irlba (Baglama, 2021) on normalized, log-transformed, and standardized counts. The principal components (PCs) are clustered using Seurat (Hao et al., 2021) using resolution 0.1 (shallow clustering) and the highest resolution that doesn’t produce any clusters containing less than 5 barcodes (deep clustering). Differential expression analysis is performed using presto (Korsunsky, 2023) between each deep cluster and all other barcodes that are not in the same shallow cluster as the deep cluster. To filter on expression-based metrics, valiDrops evaluates the top ten most significantly enriched marker genes for each deep cluster. To pass filtering, none of these genes can be expressed in at least 1% fewer barcodes in the deep cluster compared to barcodes not in the deep cluster, and they must on average be expressed in more than 30% of the barcodes in the deep cluster, less than 70% of the barcodes not in the deep cluster, and at least in 20% more of the barcodes in the deep cluster compared to barcodes not in the deep cluster. These filters remove deep clusters that are enriched for genes that are ubiquitously expressed. Next, valiDrops evaluates the total number of differentially expressed genes and their significance levels. To pass filtering, at least 1% of the tested genes for a deep cluster must be significant and the most significant gene must pass a data-adaptive threshold that is based on the relationship between the FDR-corrected P-value of the maximally significant gene in a cluster and the average difference in percent of cells expressing markers genes between the target cluster and barcodes not in the same low-resolution cluster. These filters remove deep clusters that have weak or no enrichment of specifically expressed genes. Finally, deep clusters, which have a mitochondrial or ribosomal fraction higher than three times S_n_ above the median fraction across all clusters are removed.

The fourth stage in valiDrops is to predict dead cells. Firstly, valiDrops arcsine-transforms the proportion of UMIs assigned to ribosomal genes (R) and to protein-coding genes (C) and rescales the transformed values to a range between 0 and 1 by dividing by half pi. The log transformed total UMI count (U), and the log total number of features (F) are then centered, and an initial score is calculated using the following function:

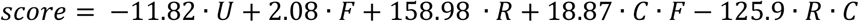

Next, initial labels are created by a data-adaptive threshold, which is defined as the first knee-point in the empirical distribution function of scores between the 0% quantile and an upper quantile of 10%. Barcodes with scores below the threshold are labeled as putative dead cells, and barcodes with scores above the threshold are labeled as putative live cells. If no barcodes are labeled as putative dead by this approach, the upper quantile is increased in steps of 10%. If less than 3 barcodes, or more than 10% of barcodes, are labeled as putative dead cells, it is likely there are no truly dead cells or that the score is likely not well-calibrated for the dataset, and label optimization is halted. If no barcodes that passed quality control, are labeled as putative dead, the barcode with the high score is temporarily relabeled as passing quality control. Next, the top 100 PCs are calculated using irlba based on the top 2,000 HVGs found using scry (Street, 2020). Next, labels are iteratively refined over maximally ten epochs. Each epoch involves first selecting features using Kendall’s tau rank correlation coefficients between the PCs and the labels and next, randomly selecting labels based on their class probability from the previous epoch (initialized with equal probability), introducing random noise by jittering the features and fitting a ridge regression model using glmnet (Friedman et al., 2010) using 5-fold cross-validation on the noised features and sampled labels. To increase robustness and assess the quality of the fit, ten independent runs of label refinement are performed and only barcodes that have the same label in at least eight runs are labeled and runs are failed if too many barcodes are not labeled.

### Benchmarking quality filtering

#### Datasets

The following public datasets were used:

1. Single-cell RNA-seq in human PBMCs (10X, 2017, 2018, 2019, 2020, 2021b): Unfiltered count matrices were downloaded from 10X Genomics.
2. Single-cell RNA-seq in human lung (Reyfman et al., 2019): Unfiltered count matrices from healthy donors were downloaded from NCBI Gene Expression Omnibus (GEO) under accession number GSE122960
3. Single-cell RNA-seq in human islets of Langerhans (Xin et al., 2018): Raw data was downloaded from the European Nucleotide Archive (ENA) under accession number GSE114297. Count matrices were generated using STARsolo (Kaminow et al., 2021), the human genome version hg38, and Ensemble gene annotations.
4. Single-nucleus RNA-seq in human PBMCs (10X, 2021a, c, d, e, f): Unfiltered count matrices were downloaded from 10X Genomics.
5. Single-nucleus RNA-seq in human brain (Pineda et al., 2021): Raw data from healthy donors was downloaded from GEO under accession number GSE174332. Count matrices were generated using STARsolo (Kaminow et al., 2021), the human genome version hg38, and Ensemble gene annotations.

#### Processing

For all datasets, all methods were run with default parameters. For EmptyDrops (Lun et al., 2019) and scCB2 (Ni et al., 2020) barcodes with an FDR ≤ 0.01 were retained. For DIEM (Alvarez et al., 2020), calls were made using debris clusters and barcodes labeled with Clean were retained. For EmptyDrops, scCB2 and EmptyNN barcodes were also filtered on the fraction of UMIs derived from mitochondrially-encoded genes using miQC (Hippen et al., 2021) for single-cell RNA-seq datasets and by removing barcodes 3 times the median absolute deviation (MAD) above the median mitochondrial fraction for single-nucleus RNA-seq, which corresponds to a strict filtering regime (Germain et al., 2020).

#### Metrics

Labels were transferred to barcoding passing quality control using Azimuth (Hao et al., 2021) and associated reference atlases and labels (PBMC mapped to PBMC reference: level 1; celltype.l1 labels, level 2; celltype.2, Islets mapped to Pancreas reference: level 1 and 2; annotation.l1, Brain mapped to Motor Cortex reference: level 1; subclass, level 2; cluster, and Lung mapped to Lung v2 (HCLA): level 1; ann_level 3, level 2; ann_fin-est_level). For each dataset, a total of nine metrics were calculated based on level 2 labels unless otherwise indicated:

1. The average mapping score is calculated by Azimuth (Hao et al., 2021) during label transfer. Higher scores indicate better label transfer; methods were ranked in decreasing order.
2. The average label score for level 2 labels is calculated by Azimuth during label transfer. Higher scores indicate better label transfer; methods were ranked in decreasing order.
3. The fraction of labels that have the correct label hierarchy, which is defined as barcodes where the transferred level 1 and level 2 labels are child-parent labels in the reference atlas. Higher fractions indicate more consistent labeling across granularities; methods were ranked in decreasing order.
4. The entropy of the top 200 most highly expressed genes in each label class was calculated using BioQC (Zhang et al., 2017). Lower entropy indicates lower heterogeneity; methods were ranked in increasing order.
5. Median local inverse Simpson’s index (LISI) (Korsunsky et al., 2019) across transferred labels in the first 10 PCs calculated from 2,000 HVGs using Seurat (Hao et al., 2021). Lower LISI indicates lower mixing of cell type labels; methods were ranked in increasing order.
6. The average silhouette width across transferred cell type labels using Euclidean distances in the first 10 PCs was calculated from 2,000 HVGs using Seurat (Hao et al., 2021). Higher silhouette width indicates a better separation of cell types; methods were ranked in decreasing order.
7. Adjusted Rand Index (ARI) was calculated using cluster labels obtained using Louvain clustering using Seurat (Hao et al., 2021) and the transferred labels. Clustering was performed with resolutions between 0.1 and 2.0 in steps of 0.1. The maximum ARI was used. High values indicate better consistency between the obtained clusters and the cell type labels; methods were ranked in decreasing order.
8. Normalized mutual information (NMI) was calculated using the same strategy as for ARI. High values indicate better consistency between the obtained clusters and the cell type labels; methods were ranked in decreasing order.
9. V-measure was calculated using the same strategy as for ARI. High values indicate better consistency between the obtained clusters and the cell type labels; methods were ranked in decreasing order.

An overall rank was calculated by calculating the average rank across all nine metrics and ranking the average in increasing order. A rank for label mapping was calculated using the first 3 metrics (mapping score, label score, and label hierarchy), a rank for expression similarity corresponds to ranks of the expression entropy metrics, a rank for label cohesion was calculated using the median LISI and the average silhouette width and a rank for label clusterability was calculated using the ARI, NMI, and V-measure. Across all ranks, rank one corresponds to the lowest average across the ranked metrics and indicates the best performance.

For benchmarking after integration, each sample was quality controlled individually, labeled using Azimuth (Hao et al., 2021), and integrated using the RPCA method in Seurat (Hao et al., 2021) with default parameters. In the integrated space, we calculated twelve metrics associated with the quality of integration, the extent of batch effects, and the consistency of biological signals using level 2 labels:

1. For each label class, the fraction of marker genes, defined as differentially expressed genes (FDR ≤ 0.05, fold change ≥ 1, AUC ≥ 0.7) in a class label compared to all other barcodes, which were detected in at least 3 samples. Higher fractions indicate higher biological consistency between datasets; methods were ranked in decreasing order.
2. The median Spearman’s rank coefficient of correlation between pseudo-bulk expression levels per class label per dataset. Higher correlations indicate higher biological between datasets; methods were ranked in decreasing order.
3. LISI (Korsunsky et al., 2019) across transferred labels in the first 10 PCs in the integrated space calculated from 2,000 HVGs using Seurat (Hao et al., 2021). Lower LISI indicates lower mixing of cell type labels; methods were ranked in increasing order.
4. LISI (Korsunsky et al., 2019) across dataset labels in the first 10 PCs in the integrated space calculated from 2,000 HVGs using Seurat (Hao et al., 2021). Higher LISI indicates better mixing of batches; methods were ranked in decreasing order.
5. The average silhouette width across transferred type labels using Euclidean distances in the first 10 PCs in the integrated space was calculated from 2,000 HVGs using Seurat (Hao et al., 2021). Higher silhouette width indicates a better separation of cell types; methods were ranked in decreasing order.
6. The average silhouette width across transferred dataset labels using Euclidean distances in the first 10 PCs in the integrated space was calculated from 2,000 HVGs using Seurat (Hao et al., 2021). Lower silhouette width indicates less separation of batches; methods were ranked in increasing order.
7. ARI was calculated using cluster labels obtained using Louvain clustering using Seurat (Hao et al., 2021) and the transferred labels. Clustering was performed with resolutions between 0.1 and 2.0 in steps of 0.1. The maximum ARI was used. High values indicate better consistency between the obtained clusters and the cell type labels; methods were ranked in decreasing order.
8. ARI was calculated using cluster labels and dataset labels performed as metric seven. Low values indicate better batch integration; methods were ranked in increasing order.
9. NMI was calculated using cluster labels and transferred labels performed as metric seven. High values indicate better consistency between the obtained clusters and the cell type labels; methods were ranked in decreasing order.
10. NMI was calculated using cluster labels and dataset labels performed as metric seven. Low values indicate better batch integration; methods were ranked in increasing order.
11. V-measure was calculated using cluster labels and transferred labels performed as metric seven. High values indicate better consistency between the obtained clusters and the cell type labels; methods were ranked in decreasing order.
12. V-measure was calculated using cluster labels and dataset labels performed as metric seven. Low values indicate better batch integration; methods were ranked in increasing order.

An overall rank was calculated by calculating the average rank across all twelve metrics and ranking the average in increasing order. A rank for expression similarity was calculated based on the conservation of marker genes and Spearman’s correlation coefficient, a rank for label cohesion was calculated using the median LISI across labels and the average silhouette width for labels, a rank for label clusterability was calculated based on the ARI, NMI and V-measure for clustering labels, a rank for batch cohesion was calculated using the median LISI across batches and the average silhouette width for batches, and a rank for batch clusterability was calculated using the ARI, NMI, and V-measure for clustering batches. Across all ranks, rank one corresponds to the lowest average across the ranked metrics and indicates the best performance.

### Benchmarking prediction of dead cells

For the O’Flanagan datasets (O’Flanagan et al., 2019), preprocessed data for sorted dead and sorted live cells were downloaded from Zenodo under DOI 10.5281/zenodo.3407791. For the Ordoñez-Rueda datasets (Ordonez-Rueda et al., 2020), raw sequencing data from 10X Genomics runs were downloaded from ENA under accession number PRJEB33078 and aligned to the human genome version hg38 using STARsolo (Kaminow et al., 2021).

Subsequently, sorted dead and sorted live cells were quality filtered using valiDrops with modifications. In the O’Flanagan datasets, the initial step was skipped as the datasets had already been filtered. Furthermore, in both the O’Flanagan datasets (O’Flanagan et al., 2019) and the Ordoñez-Rueda datasets (Ordonez-Rueda et al., 2020) a common threshold for mitochondrial filtering was calculated using the live cells and applied to the dead cells. After filtering, the datasets were combined, and dead cells were identified using valiDrops with default parameters. Barcodes that pass quality control were embedded in UMAP space (left; true labels, right; predicted labels) using Seurat (Hao et al., 2021) based on 20 PCs calculated from 500 HVGs. The quality of classification was evaluated using Matthew’s correlation coefficient (MCC), which was calculated for the initial labels, for a baseline model using logistic regression on the initial labels, and for optimization of the initial labels using valiDrops. For sensitivity analysis, we used a stratified subsampling approach. Barcodes were grouped into ten groups based on the score used as the basis for the initial labels and between 10% and 50% of truly dead barcodes were randomly selected and removed in each group. Each subsampling was repeated twenty times.

Two datasets that had not been analyzed as part of the creation of the method, were used to validate the method. First, an unsorted sample from the Ordoñez-Rueda datasets (Ordonez-Rueda et al., 2020) was quality-filtered using valiDrops with default parameters. Barcodes that pass quality control were combined with barcodes from the sorted live and sorted dead cells. The first 10 PCs based on 100 HVGs were calculated using Seurat (Hao et al., 2021) and subsequently, used for nearest neighbor analysis using RANN. Transcriptomic similarity on pseudo-bulk expression levels was measured by the normalized root-mean-square error (RMSE) on the top 100 most highly expressed genes. The RMSE was normalized by the interquartile range in the reference samples. Second, raw sequencing data from public single-cell RNA-seq data from the spleen, the lungs, and the esophagus either immediately processed (fresh) or stored for 72 hours prior to processing (Madissoon et al., 2019) were downloaded from ENA under accession number PRJEB31843, aligned to the human genome version hg38 using STARsolo (Kaminow et al., 2021) and analyzed using valiDrops with default parameters. Initial cell type labels were transferred from annotations by the original authors. Barcodes that passed quality filtering in valiDrops, but not in the original analysis, had no labels. For these barcodes, labels were labeled using majority voting in a kNN classifier using the ten nearest neighbors in the 20 first PCs calculated from 2,000 HVGs. Ties were broken by selecting the label with the smallest median Euclidean distance. Dead cell labels were only used for runs marked as successful by valiDrops. All barcodes in unsuccessful runs were labeled as live. Transcriptomic similarity on pseudo-bulk expression levels was measured per cell type by the normalized root-mean-square error on the top 100 most highly expressed genes on subsamples of either fresh cells or cells predicted to be either live or dead after 72 hours of storage. The root-mean-square error was normalized by the interquartile range in the fresh cells. For each cell type, we randomly selected (with replacement) to same number of fresh cells, predicted live after 72 hours, and predicted dead after 72 hours as there were predicted dead cells to reduce biases from differences in group size.

## Acknowledgements

This work was supported by grants from the Novo Nordisk Foundation (NNF21SA0072102 and NNF21OC0068929), as well as the Danish National Research Foundation (DNRF grant No. 141) to the Center for Functional Genomics and Tissue Plasticity (ATLAS). Computation for this project was performed using the UCloud interactive HPC system, which is managed by the eScience Center at the University of Southern Denmark. We thank all the members of the Madsen group and our colleagues in ATLAS for comments and fruitful discussions that improved the manuscript.

## Author contributions

Conceptualization: J.G.S.M. Software and methodology: J.G.S.M. and G.K. Formal analysis, investigation, and validation: J.G.S.M. and G.K. Data curation and visualization: J.G.S.M. Writing: J.G.S.M. and G.K. Resources and funding acquisition: J.G.S.M. Supervision and project administration: J.G.S.M.

